# Lipid profiles and body mass index of young students in Jordan

**DOI:** 10.1101/042697

**Authors:** Balasim Rasheed Habeeb Alquraishi, Eman Rababah

## Abstract

**Objective:** Toanalyze the lipid profile in a population of young university students in relation with their BMI.

**Methods:** This study assessed 96 students with age between 18 and 31 years old who were classified according to their sex and their body mass index (BMI). The fastingserum submitted to lipid profile analysis, including serum level of total cholesterol (TC), High(HDL-c), triglycerides (TG)were measured by using enzymatic methodandNon HDLD and a fraction of cholesterol of low (LDL-c) which calculated mathematically besides of life habits and atherogenic data.

**Results:** The mean BMI (Kg/cm2) in male (27.75±5.76) is higher than the mean BMI in female (23.17±2.78), P value (0.0001). The mean total concentration of cholesterol, LDL-c and Non HDL (mg/dl) in males (165.88±32.20, 85.00±39.94, 105.09±34.22) respectively is less than in females (194.27±52.04, 125.32±50.39, 139.14±51.35) correspondingly. The mean total concentration of HDL-c and TG (61.97±13.29, 94.80±53.65) respectively are higher in male than those in female (54.57±13.14, 71.75±35.51) correspondingly. Lipid indices, total cholesterol/HDL, LDL/HDL and Non HDL/HDL in male (2.91±1.02, 1.50±0.86, 1.87±0.99) respectively are less than those in female (3.73±1.24, 2.47±1.24, 2.75±1.25).

**Conclusions:** The obesity of young males (25.00%) is more prevalence than the obesity of young females (2.28%). The risk concentration of total cholesterol,LDL-c and Non HDLand the lipid indices (Total cholesterol/HDL,LDL/HDL and Non HDL/HDL) are higher in females than males and this indicate that the young females have more risk to develop cardiac problems in older ages.

## Introduction

Dyslipidemia or hyperlipidemia attributed to bad life style and absence of physical activities is now considered as one important factor in developing the cardiovascular diseases [1][2][3][4][5][6].

This age category (20-30 years old) is important to predict cardiovascular risk factors in the next 20-30 years of life[1][7][8][9][10].Bad habits like fast food intake, lack of exercise that enable burn out the excess lipid in blood, with other Familial factors may lead to a gradual deposit of bad lipid (LDL-C, total serum cholesterol and non HDL-C group)[1][7][11].Many studies indicate that there is a strong relationship between overweight people (obesity) and high levels of cholesterol, TC, LDL-c, and a decreased level of HDL-c[1][3][4][11][12][13][14][15][16][17].Measuring the lipid profile (total cholesterol - TC, HDL-c, TG, LDL-c and Non HDL) and the body mass index are considered to correlate their importance in deciding the risk factors of atherosclerosis [2][3][14][18][19][16][20][21].

## Methods

### Studying group

This study was performed on 96 Petra University students 18-30 years old, regardless of sex. Information about Smoking, Alcohol intake, diabetes mellitus, cardiovascular problems and family history of hypertension, diabetes mellitus, cardiovascular diseases was correlated by self administered questionnaires.

### Measurements

Blood samples withdrawn after an overnight fasting (12 h) in young people, Samples were then subjected for centrifugation to obtain serum of at least 2 ml and stored immediately in deep freeze of −17Cº for later assessment of lipid profile including total cholesterol (TC), Triglyceride(TG-CH),High density lipoprotein(HDL-c) were measured by enzymatic method uzing commercial kits (Anaheim, CA 92807,1-800-222-9880), Low density lipoprotein(LDL-c) was calculated by(Friedwald equation[22] and Non HDL-cholesterol was calculated by subtraction of HDL from total cholesterol[23].Height measured to the least centimeter, body weight measured and BMI calculated as the weight (kg) divided by the height squared (m^2^),obesity defines as BMI ≥30 Kg/meter square, Data subjected to statical analysis using ANOVA at the significance level of P ≤0.05.

## Results

The metabolism of lipid of an overnight fasting young adult compared with two important indices body mass index (BMI) and Non-HDL index is summarized in table.1

**Table (1):** Metabolisim of lipids of an overnight fasting young adults males and females with two important indices body mass index (BMI) and Non-HDL index.SD(standerd deviation)

Anthropometric and metabolic parameters according to gender
| Male |  |  | Female |  |  |
| --- | --- | --- | --- | --- | --- |
| | Mean ±SD | % | Mean ±SD | % | P |
| **Body Mass Index (Kg/Cm<sup>2</sup>)** | 27.75 ±5.76 | | 23.17 ±2.78 | | 0.0001 |
| **18.5-24.99** |  | 34.09 |  | 81.81 |  |
| **25-29.99** |  | 40.90 |  | 15.9 |  |
| **$\geq 30$** |  | 25.00 |  | 2.28 |  |
| **Total cholesterol (mg/dl)** | 165.88± 32.20 | | 194.27± 52.04 | | 0.003 |
| **$\leq 200$** |  | 74.41 |  | 54.54 |  |
| **200-239** |  | 25.59 |  | 45.46 |  |
| **LDL-c (mg/dl)** | 85.00± 39.94 | | 125.32± 50.39 | | 0.0001 |
| **$\leq 100$** |  | 62.79 |  | 31.81 |  |
| **100-129** |  | 23.26 |  | 25.00 |  |
| **130-159** |  | 13.95 |  | 43.18 | 0.005 |
| **HDL-c (mg/dl)** | 61.97± 13.29 | | 54.57± 13.14 | | |
| **$\leq 40$** |  | 4.65 |  | 9.09 |  |
| **$\geq 40$** |  | 95.35 |  | 90.90 |  |
| **Triglycerides (TG)(mg/dl)** | 94.80± 53.65 | | 71.75± 35.51 | | 0.02 |
| **$\leq 150$** |  | 88.37 |  | 95.46 |  |
| **150-199** |  | 11.63 |  | 4.54 |  |
| **NonHDL (mg/dl)** | 105.09± 34.22 | | 139.14 ± 51.35 | | 0.0001 |
| **$\leq 130$** |  | 75.00 |  | 50.00 |  |
| **$\leq 160$** |  | 13.64 |  | 20.46 |  |
| **$\leq 190$** |  | 11.36 |  | 29.55 |  |

### Anthropometric and metabolic parameters according to gender

The mean BMI (Kg/cm^2^) in males (27.75±5.76) is higher than mean BMI in females(23.17±2.78), P value (0.0001).

34.09% of males and 81.81% of females have normal weight (BMI 18.5-24.99), 40.90% of males and 15.9% of females are overweight (BMI 25-29.99) and 25% of males and 2.28 of females are obese (BMI ≥30)

The mean Total concentration of cholesterol (mg/dl) in males (165.88 ± 32.20) is less than the mean total cholesterol in females (194.27±52.04),P value (0.003). Male and female values of cholesterol ≤ 200 are (74.41%,54.54%) respectively and the male and female values of cholesterol 200-239 are (25.59%, 45.46%) individually.

The mean total concentration of LDL-c (mg/dl) in males (85.00±39.94) is less than the mean total concentration of LDL-c in females (125.32 ±50.39),p value (0.0001).

62.79% of males and 31.81% of females have LDL-c (≤100 mg/dl), 23.26% of males and 25.00% of females have LDL-c (100-129 mg/dl) and 13.95% of males and 43.18% of females have LDL-c (130-159 mg/dl).

The mean total concentration of HDL-c (mg/dl) in males (61.97±13.29) is higher than the mean total concentration of HDL-c in females (54.57±13.14), P value (0.005)

4.65% of males and 9.09% of females have HDL-c (≤40 mg/dl) while 95.35% of males and 90.90 % of females have HDL-c (≥40 mg/dl).

The mean total concentration of triglycerides (mg/dl) in males (94.80±53.65) is higher than the mean total concentration of triglycerides in females (71.75±35.51), P value (0.02)

88.37% of males and 95.46% of females have triglycerides (≤150 mg/dl) and 11.63% of males and 4.54% of females have triglycerides (150-199 mg/dl).

The mean total concentration of Non HDL (mg/dl) in males (105.09±34.22) is lower than the mean total concentration of Non HDL in females (139.14±51.35), P value (0.0001)

75% of males and 50% of females have Non HDL (≤130 mg/dl), 13.64 % of males and 20.46% of females have Non HDL (≤160 mg/dl) and 11.36% and 29.55% of females have Non HDL (≤190 mg/dl).

### lipid indices of male and female

**Table (2):** Comparisone of lipid indices between male and female groups.(M=Mean±SD)

| **Lipid Indices** | **Male**<br>**M** | **Female**<br>**M** | **P value** |
| --- | --- | --- | --- |
| Total Cholesterol/HDL | $2.91 \pm 1.02$ | $3.73 \pm 1.24$ | 0.001 |
| LDL/HDL | $1.50 \pm 0.86$ | $2.47 \pm 1.24$ | 4.39E-05 |
| NON HDL/HDL | $1.87 \pm 0.99$ | $2.75 \pm 1.25$ | 0.0005 |

The mean Total Cholestrol/HDL in males (2.91±1.02) is less than the same ratio in females (3.73±1.24), p value ≤0.001

The mean LDL/HDL in males (1.5±0.86) is less than this ratio in females (2.47±1.24),P value ≤ 4.39E-05

The mean Non HDL/HDL ratio in males (1.87±0.99) is less than the same ratio in females (2.75±1.25), P value ≤0.0005

## Discusstion

The incidence of risk factors for cardiovascular diseases in young adults especially for the university students must be given an importance. Several studies indicate that obesity is the main factor of dyslipidemia which is one of the most important etiological initiator for cardiovascular diseasesand hypertension [2][15]

The studies showed that dyslipidemia associated with obesity is characterized by high level of triglycerides, low levels of HDL, and abnormal LDL arrangement [15] [24]

Our study evaluated 96 students aged between 18 and 31 years old who were classified according to their sex and their body mass index (BMI). Their fasting serums over night (12 h) were analyzed for lipid profile, including serum levels of total cholesterol (TC), High(HDL-c), triglycerides (TG),Non HDLD and fraction of cholesterol of low (LDL-c) which calculated mathematically.

Mean values of BMI were significantly higher in male than in female (P≤0.0001) and This can be attributed to the presence of excess muscle mass in male body which is equivalent to almost twice the proportion of muscle in a female body so male BMI is higher than BMI of female due to muscles rather than the presence of fat[25].

The mean Total concentration of cholesterol (mg/dl) in male (165.88 ±32.20)is less than the mean total cholesterol in female(194.27±52.04),P value (0.003). Male and female value of cholesterol ≤ 200 are (74.41%,54.54%) respectively and the male and female values of cholesterol 200-239 are (25.59%, 45.46%). The increase in serum cholestrol in females mostly due to lack of exercise and ingestion of fatty meals which are rich with cholestrol.In spite of that females are protected by increased level of estrogen that affect plasma lipid and exert benefical effect on carbohydrate metabolism[26].

LDL-c Mean ± SD values in males (85.00± 39.94)) is less than same data in female (125.32 ± 50.39 mg/dl where P≤0.0001.In spite of that difference in mean, fractional values show some controversy as,LDL-c percentage≤100 mg/dl is more in male than female group (M 62.79% Vs. F 31.81%), LDL-c category of 100-129 mg/dl both group shows mild insignificant differences of (M 23.26% Vs. F 25.00%) but in extremely high LDL-c value show the reverse where the male value is significantly lower than that of the female (M 13.95% Vs. 43.18%) this explain why mean values show significant differences.

HDL-c in males values is significantly higher than in females (61.97±13.29 mg/dl vs. 54.57±13.14 mg/dl), P ≤0.005. Fractional values of this protective lipid indicate that males HDL-c of ≥40 mg/dl is significantly higher than that in females (M 95.35% vs. F 90.90%) while HDL-c of ≤40 mg/dl is significantly less in males than in females group (M 4.65%vs. F 9.09%).

TG-c mean values of male group are significantly higher than female group (94.80± 53.65 mg/dl and 71.75± 35.51 mg/dl) respectively P ≤0.02.This can be attributed to the high level of calories in male food intake with little sprorts and excerice.

Non-HDL male mean values are significantly less than female values (M 105.09± 34.22 mg/dl vs. F 139.14 ± 51.35mg/dl) p=0.0001.This is logical because the mean total concentration of cholestrol in male is less than in female and the mean total concentration of HDL in males is higher than in females.

Non-HDL tend to be more in females than male groups except in the (≤130 mg/dl) category where it is mildly decreased and this can be explaind by the variation of the type of food and its contents of fats.

Lipid indices: Total cholesterol/HDL, LDL/HDL and Non HDL/HDL in male (2.91±1.02,1.5±0.86,1.87±0.99) respectively are less than those in female (3.73±1.24,2.47±1.24,2.75±1.25).These results align with previous results which indicated that the mean concentration ofcholesterol, LDL-c and Non HDL in males are less than the same mean concentration in females while the mean concentration of HDL-c and TG in males are higher than females.

## Conclusion

The obesity of young males is more prevalent than the obesity of young females. The mean total concentration of cholesterol, LDL-c and Non-HDL are less in male than female while the mean total concentration of HDL-c and TG are higher in male than in female. The lipid indices (Total cholesterol /HDL, LDL/HDL Non-HDL/HDL) are less in male than female.

These results indicate that the young females may be subjected to developing cardiac problems in older age.

According to above data of this study and other investigator findings we recommend an evaluation of the link between lipid values and expecting risk factors upon the cardiovascular system.

